# mutSigMapper: an R package to map spectra to mutational signatures based on shot-noise modeling

**DOI:** 10.1101/2020.10.12.336404

**Authors:** Julián Candia

## Abstract

**Summary:** mutSigMapper aims to resolve a critical shortcoming of existing software for mutational signature analysis, namely that of finding parsimonious and biologically plausible exposures. By implementing a shot-noise-based model to generate spectral ensembles, this package addresses this gap and provides a quantitative, non-parametric assessment of statistical significance for the association between mutational signatures and observed spectra.

**Availability and implementation:** The mutSigMapper R package is available under GPLv3 license at https://github.com/juliancandia/mutSigMapper. Its documentation provides additional details and demonstrates applications to biological datasets.

## Introduction

A mutational signature is a quantitative representation of a mutagenic process in a discrete space of somatic mutation motifs. Once a set of such signatures (*compendium*) is inferred from thousands of cancer genomes and exomes, ^1,2^ or established by mutagen treatment experiments, ^3^ the mutational profile (*spectrum*) of individual cancer samples can be compared against the compendium to inform of possible etiologies, features for prognostic and biologic stratification, as well as vulnerabilities to be exploited therapeutically. Although many types of genomic alterations can serve as features of mutational signatures, most studies to date have focused on frequency profiles along the 96 trinucleotide contexts (*channels*) centered around all possible single-base substitutions (Fig. 1a). The observed spectra can thus be described as linear combinations (*exposures*) of the signatures in the compendium (Fig. 1b).

**Figure 1:**
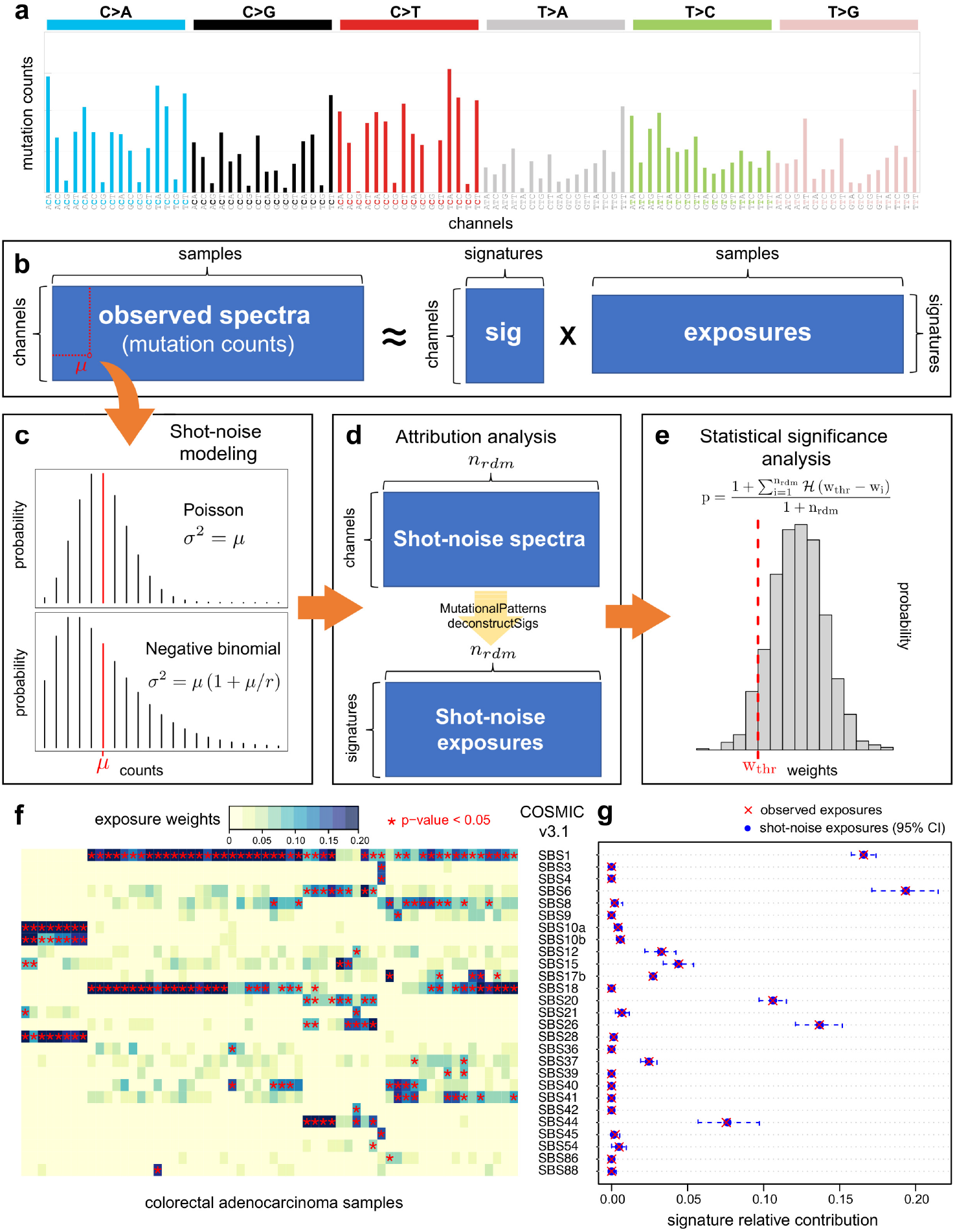
(a) Mutational profile using the conventional 96-channel representation. (b) Observed spectra expressed as the matrix product of signatures by exposures. (c) The observed mutation count of one sample in a given channel can be expanded as a Poisson or negative binomial shot-noise distribution. (d) Solving the attribution problem for each random sample, we obtain a distribution of exposures (e), whose statistical significance is assessed via non-parametric p-values as shown (where is the Heaviside function). (f) 60 WGS colorectal adenocarcinoma samples mapped to the COSMIC v3.1 compendium (only significant signatures shown). (g) Detailed results for one patient.

Based on the analysis of a large number of spectra, the “extraction and attribution” approach focuses on simultaneously finding de-novo signatures and exposures, usually implemented as non-negative matrix factorization algorithms.^4–6^ The “attribution-only” approach, in contrast, assumes an existing compendium and aims to find the exposures that optimally reconstruct the observed spectra using standard constrained linear optimization methods. ^7,8^

De-novo signature discovery is a challenging task with several known issues, among them: defining the optimal number of signatures, discovering weaker signatures in a background of stronger ones, and resolving spill-over (*bleeding*) between signatures. Moreover, shared with the “attribution-only” approach, another key issue is that of finding parsimonious and biologically plausible exposures. Indeed, adding contributions from more signatures typically improves reconstruction, even when those additional etiologies are biologically implausible. As an illustration of the latter, the reconstruction of colorectal adenocarcinoma spectra (Suppl. Fig. 35 in Alexandrov et al, 2013^1^) shows an ubiquitous contribution of COSMIC signature 6, characteristic of microsatellite instability (MSI) resulting from impaired DNA mismatch repair (MMR), although only about 15% of colorectal tumors are known to arise due to MSI/MMR deficiencies. ^9,10^ Other common examples are ultraviolet signatures in tumors with no possibility of UV exposure, and signatures of defective polymerase epsilon proofreading (which generates very high numbers of mutations) with very low mutation counts and no defect in the polymerase epsilon gene. The association between spectra and signatures is often visualized as cosine similarity heatmaps ^8,11^ that put these shortcomings in clear evidence. Here, the central question we aim to address is *how to assess, in a statistically meaningful way, the significance of the association between spectra and mutational signatures?* With mutSigMapper, we propose a framework to robustly map spectra to signatures based on shot-noise modeling.

## Method

Shot noise is the most fundamental model of discontinuous noise in continuous-time physical systems, used e.g. to model low-exposure optical phenomena in which events (the detection of photons) are statistically independent and relatively rare. ^12^ Shot noise is often modeled as a Poisson process, characterized by a probability distribution whose mean equals its variance, which is generalized by the negative binomial distribution (Fig. 1c). Analogously to low-exposure physical phenomena, the occurrence of mutagenic events along the genome is well modeled as a Poisson-like process. ^13^ Under this perspective, the observed mutation count from a sample in a given channel is a random variable that follows a shot-noise probability distribution whose mean may be set to the observed mutation count. From each sample, therefore, we can randomly generate *n_rdm_* shot-noise spectra (Fig. 1d). By using a standard “attribution-only” approach, we can then assess the contribution of each signature to each of these random spectra, thus leading to a distribution of exposure weights. Finally, by calculating an empirical p-value based on the resulting distribution, ^14^ we can hence quantitatively assess the statistical significance of the observed contribution of each signature to the sample’s mutational spectrum (Fig. 1e).

## Implementation

The main input to mutSigMapper is the spectra data frame that contains mutation counts, which can be extracted from VCF or MAF files using existing software tools. ^11,15^ The package offers built-in compendia from COSMIC and mutagen exposure resources; alternatively, the user may provide a custom signature data frame. Built-in noise model choices are Poisson and negative binomial. Non-negative least squares regression procedures (implemented via MutationalPatterns and deconstructSigs) are run for the observed spectra and for the *n_rdm_* shot-noise random spectra. Data exporting and visualization methods are provided in the package, as well as case-example datasets and detailed documentation.

## Case example

We extracted mutational spectra from 60 WGS colorectal adenocarcinomas and applied mutSigMapper to find associations with the 72 signatures from the COSMIC v3.1 compendium. By generating *n_rdm_* = 1000 realizations of Poisson noise (Fig. 1f-g), we found that Signature SBS6 is significantly associated with colorectal cancer in 13.3% of the cohort, in excellent agreement with known MSI prevalence. The ubiquitous Signature SBS1 (an endogenous mutational process initiated by spontaneous deamination of 5-methylcytosine) is found in all cancer types and in most cancer samples. Other prevalent signatures are SBS10 (polymerase epsilon exonuclease domain mutations) and SBS18 (damage by reactive oxygen species), in agreement with previously reported observations. ^2^ For step-by-step details and further discussion of the effect of mutational burden and noise models, see the accompanying vignette (Supplementary Information).

## Conclusions

Exploiting the analogy between mutagenic exposures and shot noise phenomena in optics and electronics, we propose a model to generate spectral ensembles that allow a quantitative, non-parametric assessment of statistical significance for the association between mutational signatures and observed spectra. The package mutSigMapper thus fills an important gap in the existing software for mutational signature analysis.

## Supporting information

Package Vignette

## Declarations

## Acknowledgements

The author thanks Steven G. Rozen, Arnoud Boot, and Ferran Muiños for insights shared during ISMB 2020’s Mutational Signatures tutorial lectures.

