## Supplementary material for "mutSigMapper: an R package to map spectra to mutational signatures based on shot-noise modeling": Package Vignette

### mutSigMapper Vignette

#### Table of contents


1. About
2. Installation
3. Case example: Colorectal adenocarcinoma
4. Case example: Melanoma


#### 1. About

The R package `mutSigMapper` is available under GPL-3 license at https://github.com/juliancandia/mutSigMapper.

Author and maintainer: Julián Candia <>

#### 2. Installation

To install this package:

```
if (!require("devtools")) {
    install.packages("devtools")
}
devtools::install_github("juliancandia/mutSigMapper")
```

#### 3. Case example: Colorectal adenocarcinoma

Let us analyze the PCAWG dataset consisting of 60 whole-genome-sequencing (WGS) colorectal adenocarcinoma samples (Whole Genomes Network 2019). First, we load the `mutSigMapper` package:

```
library(mutSigMapper)
```

Next, we load the `WGS_PCAWG` dataset and select the colorectal adenocarcinoma samples:

```
data(WGS_PCAWG)
COAD_PCAWG_index = which(WGS_PCAWG$sample_study[, "study"] == "ColoRect-AdenoCA")
```

Now we run `mutSigMapper` on these samples against the COSMIC v3.1 (June 2020) compendium:

```
set.seed(123)
COAD_PCAWG_v3.1 = mutSigMapper(WGS_PCAWG$spectra[, COAD_PCAWG_index], ref = "cosmic_v3.1",
    n_rdm = 1000)
```

The significant signatures present in this cohort are summarized by

```
weights = plotSpectraHeatmap(COAD_PCAWG_v3.1, signif.sig.only = T, cexRow = 0.8)
```

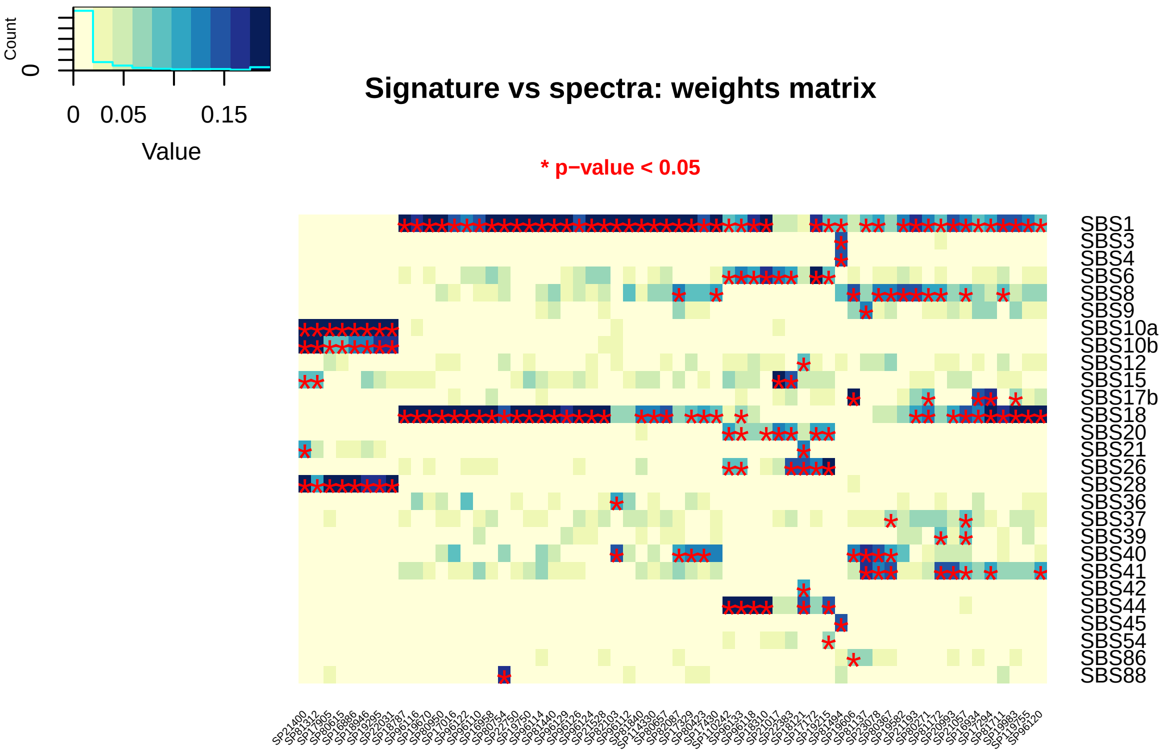


The percentage of samples associated with signature “SBS6”, which captures microsatellite instability events resulting from impaired DNA mismatch repair (MSI/MMR), is:

```
100 * sum(COAD_PCAWG_v3.1$map_pval["SBS6", ] < 0.05)/length(COAD_PCAWG_index)
```

```
## [1] 13.33333
```

This result is in excellent agreement with clinical reports for colorectal cancer (Boland and Goel 2010; Sinicrope 2010).

Let us now examine the TCGA dataset consisting of 496 whole-exome-sequencing (WES) colorectal adenocarcinoma samples:

```
data(WES_TCGA)
COAD_TCGA_index = which(WES_TCGA$sample_study[, "study"] == "ColoRect-AdenoCa")
```

Because the COSMIC v3.1 compendium is genome-based, we adjust the signatures using the `sig.bkg.adj` argument to reflect exomic background frequencies:

```
set.seed(123)
COAD_TCGA_v3.1 = mutSigMapper(WES_TCGA$spectra[, COAD_TCGA_index], ref = "cosmic_v3.1",
    sig.bkg.adj = "exome/genome", n_rdm = 1000)
```

The resulting percentage of WES samples associated with signature “SBS6” is:

```
100 * sum(COAD_TCGA_v3.1$map_pval["SBS6", ] < 0.05)/length(COAD_TCGA_index)
```

```
## [1] 1.814516
```

This smaller percentage, compared with our WGS analysis before, reflects the fact that MSI/MMR events in exonic regions are much less prevalent than in non-exonic ones (even after adjusting by background microsatellite abundance) (Kim, Laird, and Park 2013).

### 4. Case example: Melanoma

Let us now consider melanoma, the human cancer type that carries the highest burden of somatic mutations (Alexandrov et al. 2013). The PCAWG dataset contains 107 WGS melanoma samples:

```
data(WGS_PCAWG)
SKCM_PCAWG_index = which(WGS_PCAWG$sample_study[, "study"] == "Skin-Melanoma")
```

```
set.seed(123)
SKCM_PCAWG_v3.1 = mutSigMapper(WGS_PCAWG$spectra[, SKCM_PCAWG_index], ref = "cosmic_v3.1",
    n_rdm = 1000)
```

The significant signatures present in this cohort are summarized by

```
weights = plotSpectraHeatmap(SKCM_PCAWG_v3.1, signif.sig.only = T, cexRow = 0.8)
```

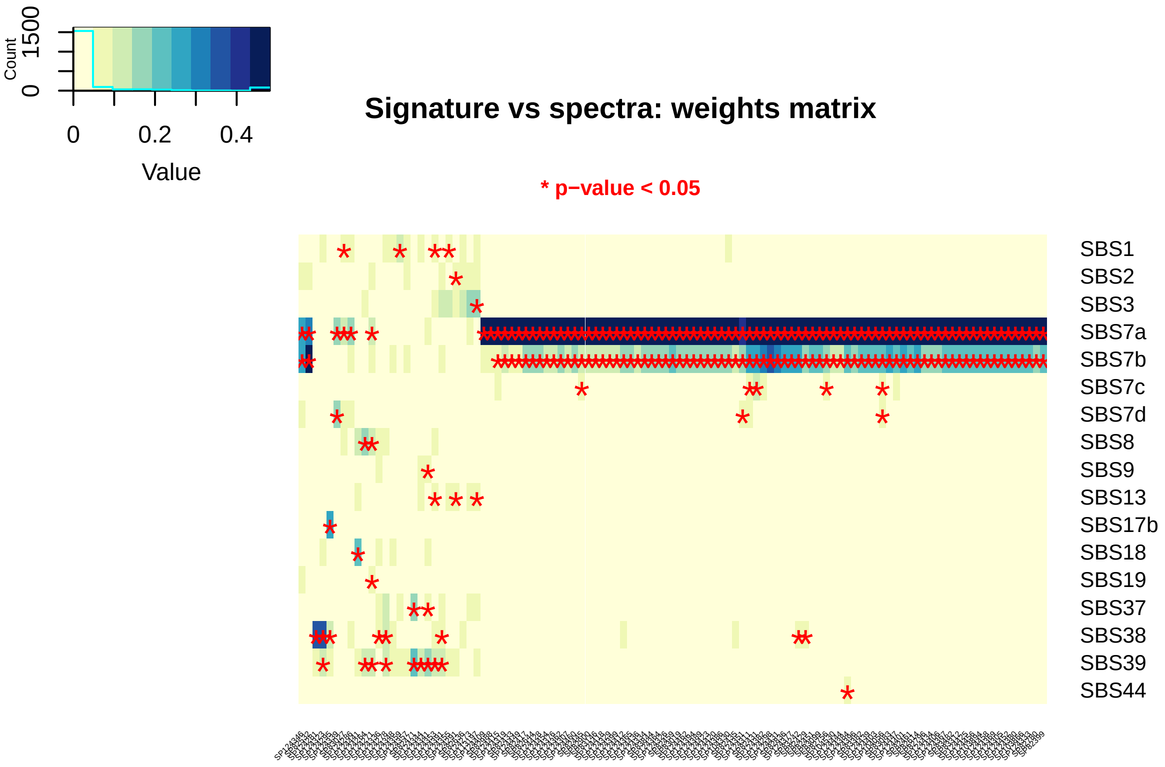


The most prevalent signatures are "SBS7" subtypes, all of them associated with UV exposure. To explore the prevalence of the two most frequent subtypes:

```
SBS7ab_prev = round(100 * table(SKCM_PCAWG_v3.1$map_pval["SBS7a", ] < 0.05, SKCM_PCAWG_v3.1$map_pval["S
    ] < 0.05)/length(SKCM_PCAWG_index), digits = 1)
rownames(SBS7ab_prev) = paste0("SBS7a:", c("no", "yes"))
colnames(SBS7ab_prev) = paste0("SBS7b:", c("no", "yes"))
SBS7ab_prev
```

```
##
##               SBS7b:no SBS7b:yes
##   SBS7a:no      18.7       0.0
##   SBS7a:yes      5.6      75.7
```

This tells us that the "SBS7a" subtype is present in 81.3% of the samples; most of these (75.7% of the total cohort) are also associated with the "SBS7b" subtype.

Spectra vs signature similarity metrics (cosine similarity, correlation, Jensen- Shannon divergence) are also available for further exploration. For instance:

```
cos_sim = plotSpectraHeatmap(SKCM_PCAWG_v3.1, signif.sig.only = T, cexRow = 0.8,
    type = "cosine")
```

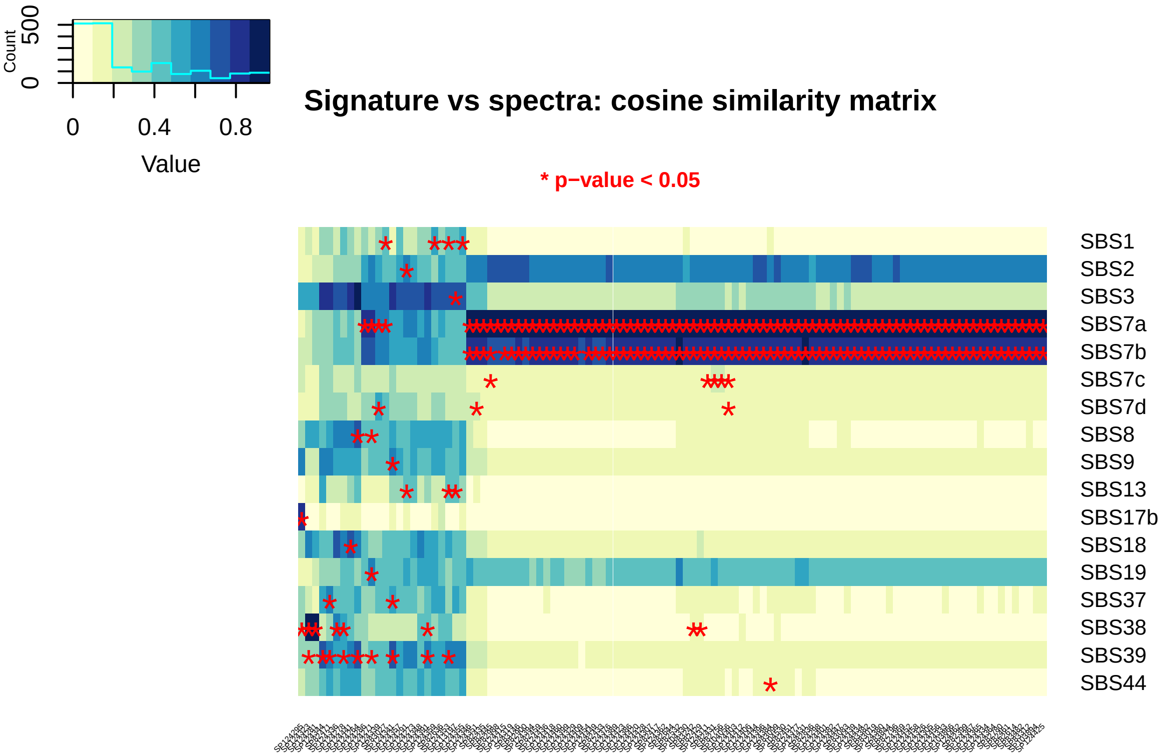


Signatures such as "SBS2", which appears similar to many of the spectra, are nonetheless not significantly associated. By examining the cosine similarity across signatures in COSMIC v3:

```
cos.sim = function(ma, mb) {
    mat = tcrossprod(ma, mb)
    t1 = sqrt(apply(ma, 1, crossprod))
    t2 = sqrt(apply(mb, 1, crossprod))
    mat/outer(t1, t2)
}
data(ref_cosmic_v3.1)
cosmic_v3.1 = t(ref_cosmic_v3.1$sig[, -c(1, 2)])
cosmic_v3.1_cos_sim = cos.sim(cosmic_v3.1, cosmic_v3.1)
```

we find that the signature most similar to "SBS2" is "SBS7a":

```
head(cosmic_v3.1_cos_sim["SBS2", order(-cosmic_v3.1_cos_sim["SBS2", ])])
```

```
##      SBS2     SBS7a     SBS30     SBS58     SBS40     SBS50
## 1.0000000 0.7142544 0.4613371 0.3869516 0.3312233 0.2891235
```

Thus, it becomes apparent that the observed similarity between "SBS2" and many of the spectra is not causal, but mediated by the UV-exposure signature "SBS7a" that `mutSigMapper` identified as significantly associated.

Let us examine in greater detail one sample that apears associated with all "SBS7" subtypes:

```
spectrum = "SP104056"
SKCM_PCAWG_v3.1$map_pval[SKCM_PCAWG_v3.1$map_pval[, spectrum] < 0.05, spectrum]
```

```
##       SBS7a       SBS7b       SBS7c       SBS7d
## 0.000999001 0.000999001 0.002997003 0.000999001
```

Notice that for subtypes "SBS7a,b,d", the empirical p-values reached the theoretical minimum imposed by $1/(n_{rdm} + 1)$ (Phipson and Smyth 2010). That is, the statistical power is limited by the number of random realizations generated. If we want to explore p-values smaller than this theoretical minimum, we need to re-run `mutSigMapper` with accordingly larger $n_{rdm}$.

We can plot the list of top-15 signatures from the Cosmic v3.1 compendium associated with this sample:

```
weights = SKCM_PCAWG_v3.1$weights[[spectrum]]
sigs = names(weights)[order(-weights)[1:15]]
plotSpectraCaterpillar(SKCM_PCAWG_v3.1, spectra.set = spectrum, sig.set = sigs, cexRow = 0.85)
```

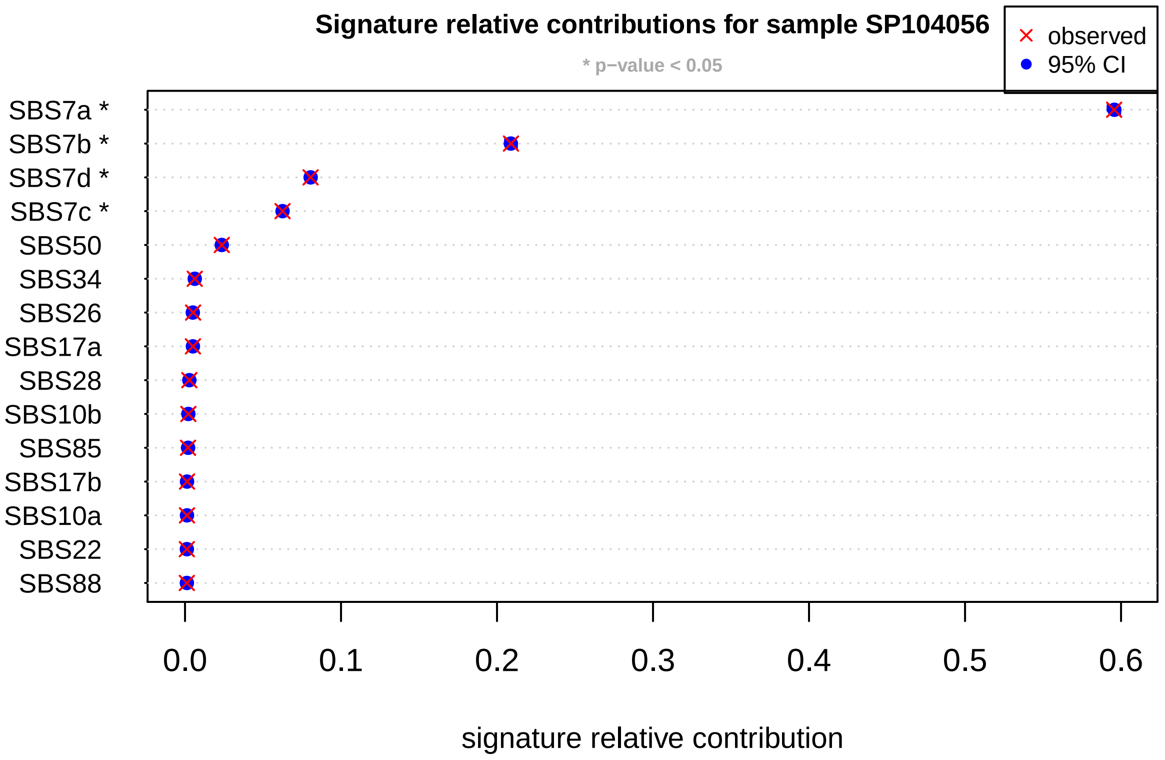


Let us examine the effect of shot noise model; we will analyze negative binomial noise with different size parameters. First, we generate `mutSigMapper` objects for different size levels:

```
size = 10^seq(-1, 4, by = 1)
n_size = length(size)
noise_effects = vector("list", n_size)
for (i_size in 1:n_size) {
    set.seed(123)
    noise_effects[[i_size]] = mutSigMapper(WGS_PCAWG$spectra[, spectrum, drop = F],
        ref = "cosmic_v3.1", noise = "neg.binom", neg.binom.size = size[i_size],
        n_rdm = 1000)
}
```

Then, we extract and merge the p-values:

```
map_pval_neg_binom = NULL
for (i_size in 1:n_size) {
    map_pval_neg_binom = cbind(map_pval_neg_binom, noise_effects[[i_size]]$map_pval)
}
```

We use this object to analyze the attribution of "SBS7" signatures across sizes:

```
target_sig = paste0("SBS7", c("a", "b", "c", "d"))
n_target = length(target_sig)
plot(range(size), range(-log10(map_pval_neg_binom[target_sig, ])), log = "x", type = "n",
    xaxt = "n", xlab = "negative binomial size parameter", ylab = "-log10(p-value)")
axis(1, labels = c(0.1, 1, 10, 100, 1000, 10000), at = size)
for (i_target in 1:n_target) {
    lines(size, -log10(map_pval_neg_binom[target_sig[i_target], ]), type = "b", col = i_target)
}
abline(h = -log10(0.05), col = "black", lty = 2)
legend(0.1, 3, c(target_sig, "p-val=0.05"), col = c(1:n_target, "black"), lty = c(rep(1,
    n_target), 2), pch = c(rep(1, n_target), NA))
```

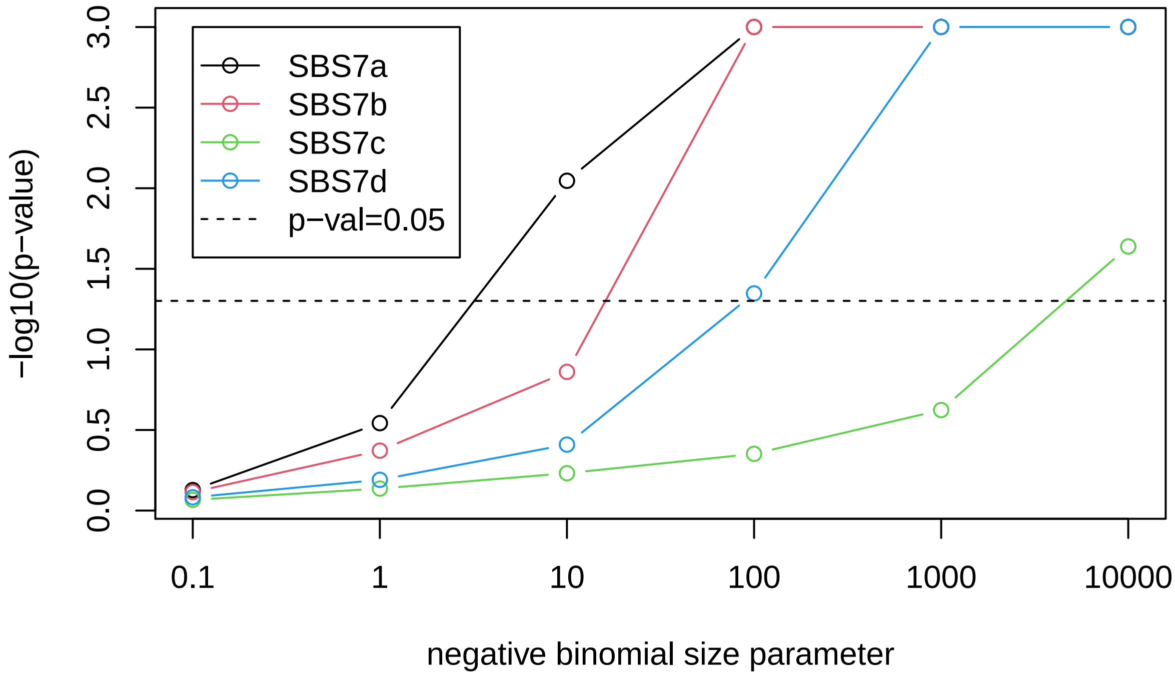


Notice that, as we increase the negative binomial size parameter, solutions increasingly resemble the results obtained with Poisson noise, as expected. The effect of negative binomial distributions on signature attribution is well illustrated by caterpillar plots generated for different values of the `size` parameter. For instance, compare `size=1`:

```
plotSpectraCaterpillar(noise_effects[[which(size == 1)]], sig.set = sigs, main.title = "Size=1")
```

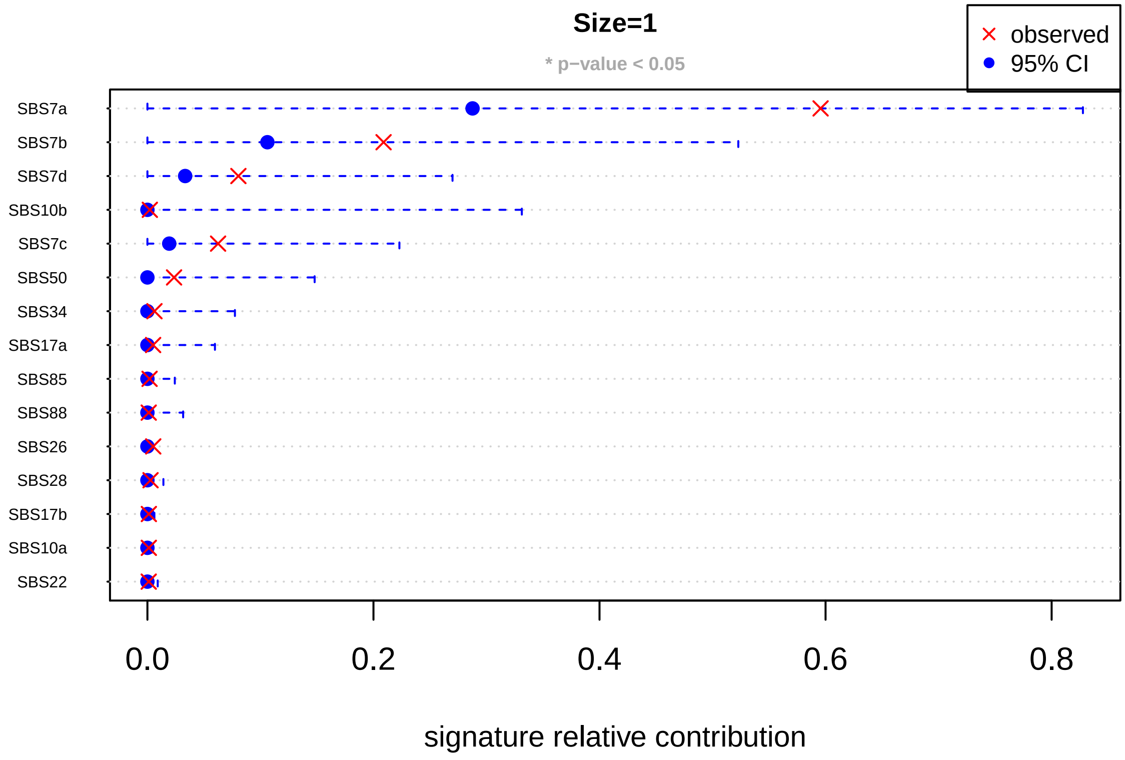


with `size=100`:

```
plotSpectraCaterpillar(noise_effects[[which(size == 100)]], sig.set = sigs, main.title = "Size=100")
```

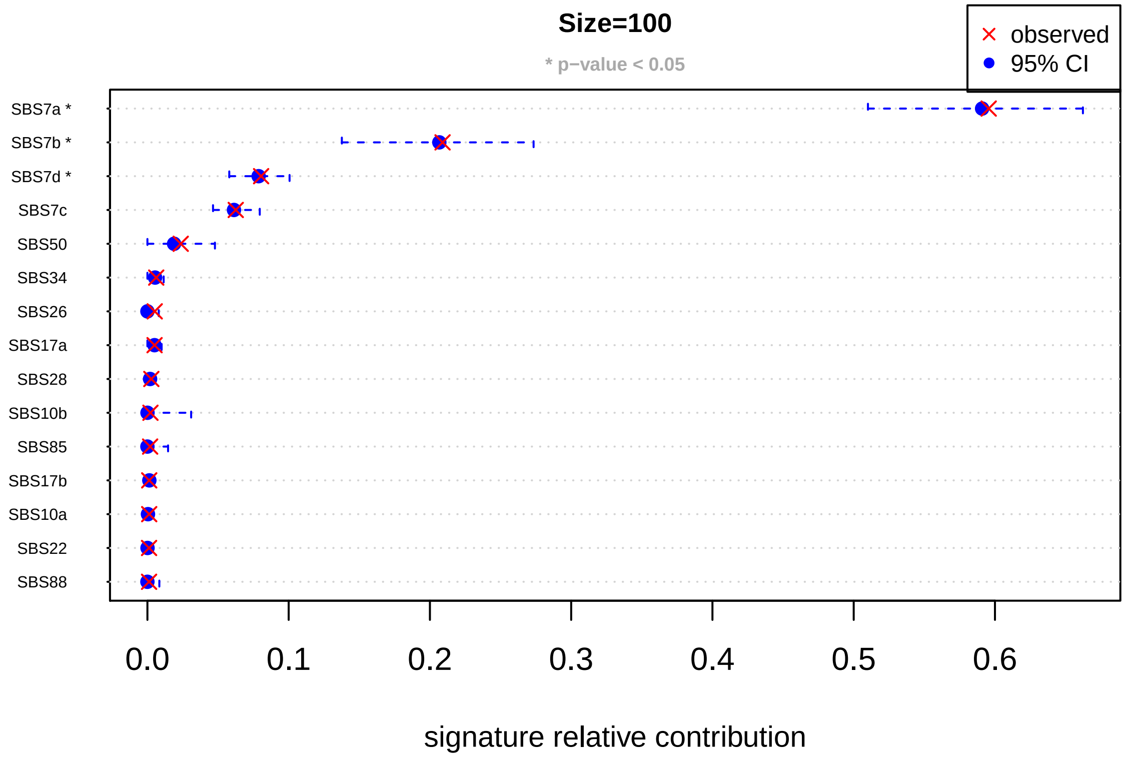
